# Silver spoon effect: early-life environment and adult survival in a primate

**DOI:** 10.64898/2026.09.23.753472

**Authors:** Martine Perret, Aude Anzeraey, Fabienne Aujard

## Abstract

Early environmental conditions can have long-lasting effects on survival and reproductive fitness. Using a 23 year-long monitoring data set of captive mouse lemur’s life history traits, we tested a relationship between maternal allocation to offspring (N = 1365) and their adulthood survival. Maternal characteristics such as age or body condition did not affect allocation to offspring whatever the litter size (1 to 3). Although birth mass depended on size and composition of the litters, mouse lemur survival was highly correlated with body mass acquired after weaning in both sexes, through potential sibling competition. Moreover, female’s reproductive success correlated with this body mass and was consistently associated with increased longevity. These findings suggest the presence of a silver spoon effect under constant captive conditions. However, cumulative effects of genetic and adulthood social conditions strongly interact to affect adult survival. Deaths related to intra- or inter-sexes aggressive social interactions may outweigh effect of early environment and therefore minimize the silver-spoon effect on captive mouse lemur’s survival. However, having a high body mass after weaning appeared to be a determinant factor in individual survival for mouse lemurs.

## INTRODUCTION

A growing number of studies on vertebrates showed that early life conditions could have long-term consequences on individual fitness-related traits, mainly reproductive success and survival [1-4]. The concept of the ‘silver-spoon effect” [5] predicts that individuals experiencing favourable early-life environment might benefit from lifelong positive effects. A greater energy acquisition during development would allow individual to invest simultaneously in several traits, consistent with expectations for trade-offs between reproduction and survival. For example, growing quickly to reach a large size would give advantage to reproduce earlier and potentially to live longer. Early life conditions may be positive from high level of maternal care or negative from sibling competition known to influence late-life physiological and behavioural individual phenotype [6-8]. Maternal care depends of numerous factors such as body condition, parity, age and resources available for allocation to offspring, all of these factors being themselves dependent of environmental factors.

In wild animals, mostly in females, it is predicted that individuals exposed to poor early-life conditions may respond by different strategies to maximise their fitness: delay in reproduction associated with lower senescence rate or early life reproduction at the cost of survival [9,10]. Several studies in birds and mammals revealed that poor early resource availability, early high population density, poor natal habitat quality or early predation risk lead to a lower quality of offspring and then, a decrease in life–time reproductive success during adulthood. This last effect can occur with or without effect on survival owing the sex-specific complexity of ageing patterns [1,2, 11-22]. Moreover, these effects may differ between sexes depending on selection pressure [12, 13]. On the opposite, favourable early resource conditions provide long-lasting “silver spoon” effects [23-25]. Lastly, changes in behaviours and cognitive performances were also described in adults having experienced either early adversity [26-28] or high favorable conditions [24].

Such effects of early unfavourable conditions on fitness and survival risk have been reported in several species of primates [29-33]. Likewise, studies on human populations demonstrated the significant influence of early unfavourable conditions (famine, war) on morbidity and mortality in adulthood [9, 34-38].

In primates, the complexity of social structures and the role of both parents for offspring care render difficult to elucidate the role of maternal conditions on late-life offspring fitness. By contrast, the grey mouse lemur, a small Malagasy primate (*Microcebus murinus*), offers several practical advantages for testing relationship between maternal allocation to offspring and adulthood survival. In this species, which longevity may reach up to 13 years [39], females may give birth to 1 to 3 infants per year, nursing them without paternal care. The constant environmental conditions of captivity allow testing long-term effects of maternal allocation without any other confounding factors. Previous studies have shown that littermates have long-term effects on reproductive fitness in both sexes. Reproductive fitness is reduced in females born with a male litter mate, whereas males born in mixed sex litters were more competitive during sexual encounters [40,41]. These results suggested a potential long-term effect on survival through early conditions experienced by offspring.

Using a 23 year-long monitoring data of mouse lemur’s life-history traits, we focused on the effect of the mother condition and on its allocation during nursing, considering the size and the composition of the litter in which offspring were born. In captivity, where conditions are constant, we predicted that offspring receiving high maternal care have greater high body mass at weaning and would have a better adulthood survival. Finally, we asked for a potential sex-specific “silver-spoon” effect, if any.

## MATERIAL and METHODS

### Biological model

The grey mouse lemur is a small primate endemic of Madagascar. Mouse lemurs are strict long day breeders and exhibit seasonal biological rhythms in response to both dry winter and wet summer seasons occurring in the wild. During the winter season, resources are scarce and animals use energy-saving mechanisms (fattening, decreased activities and torpor). Reproduction is highly seasonal with females exhibiting marked oestrous synchrony and males entering sexual competition.

The breeding colony of mouse lemurs was established 60 years ago from a stock originally caught near the southern western coast of Madagascar nearly sixty years ago. For several decades, mouse lemurs have been studied as a model for investigating various aspects of biology and aging processes. Captive conditions were maintained constant with respect to ambient temperature (24–26°C) and hygrometry (55–60%). Animals were fed *ad libitum* on a standardized diet, including fresh fruits, a homemade milky mixture (19.3% proteins, 17.2% lipids and 63.5% carbohydrates) and mealworms. To ensure seasonal reproductive rhythms, animals were routinely exposed to an artificial light photoperiodic cycle consisting of 6 months of long photoperiod (LD = 14 h of light/day, summer-like) followed by 6 months of short photoperiod (SD = 10 h of light/day, winter-like). At the time of LD exposure, groups composed of 2-3 unrelated males and 1 to 3 females were randomly constituted. Immediately, males entered competition for priority access to oestrous females, leading to a hierarchy mostly depending on aggressive interactions [42]. After mating, females were isolated for pregnancy and lactation. Females give birth to 1 to3 young per litter, exceptionally 4, after a gestation period lasting approximately 2 months. They nurse infants for 30-40 days without paternal care. When 3 months-old, offspring were removed from their mother and housed in 4-5 single-sex groups. Outside the mating period, males and females were separately housed in single sex groups (2-4 individuals).

### Data analysis

To investigate the relationship between effect of maternal allocation and survivorship of offspring, we analysed longitudinal data recorded over 23 years (2000 to 2023 included) on 2800 infants (1406 females, 1394 males) from birth to death. The studied population originated from 1314 litters issued from 854 different mothers. Female mouse lemur can reproduce from the first breeding season until death [43-,44]. Neonatal mortality (N = 398, 14% of births), as well as offspring that died before one year (N = 82, mean survival: 0.53± 0.03 yrs), were not included. Censored adults either died after experimental purposes (N = 270), were transferred in another breeding colonies (N = 297) or were still alive at the moment of the data analysis (N = 388). Thus, our analyses were restricted to individuals born between 2000 and 2023 that had completed natural lifespan, (N = 1365, 717 females and 648 males, mean survival: 5.55 ± 0.06 yrs).

Several parameters were selected.

For each offspring, maternal parameters included age at conception, parity, body mass at oestrus, an indicator of resources that the mother could allow to her offspring (BM*oe*), and gestation length.

For offspring, we recorded body mass at birth (BM*birth*), at 30 days i.e mass gain during lactation (BM*30d*), and 2 months after birth, i.e after weaning (BM*60d*). Early growth rate was estimated by the slope of regression of body mass from birth to 30 days old. This represents an estimation of maternal allocation to each individual early in life. Maternal allocation and sibling competition depended on the size of the litter in which offspring were raised: from 1 to 3 offspring (no litters of 4 were present). The composition of the litter was also considered: male litters (M, MM, MMM), female litters (F, FF FFF) and mixed-sex litters (MF, MMF, MFF).

The reproductive success concerned only females. All females had the opportunity to breed at least once during their life leading to 511 reproductive successes (offspring raised until weaning) and 206 breeding failures (unsuccessful fecundation, abortion or dead offspring). The lack of genetic analyses for many males did not allow determination of reproductive success of all males.

Finally, age and photoperiodic regimen at death were included and, if available, cause of death determined by visual examination and autopsy.

### Statistics

Data are presented as mean ± SEM. Statistical analyses included Cox proportional hazards model for survival curves, multi-way analyses of variance using or not litter size a covariate for relationships between parameters of offspring and litter’s composition, maternal characteristics and adulthood survival.

G tests were used to test distributions. Multiple pairwise comparisons were made using Tukey’s post hoc test. In addition, relationships between the different factors were tested using linear regression analyses. All statistical analyses were conducted using Systat Software. To avoid overloading the text, statistical details were not systematically given for insignificant probabilities.

### Ethics Statement

All the results in this study did not correspond to experimental procedures but are issued from the exploitation of data collected in the mouse lemur’s captive population. We adhered to the Guidelines for the Treatment of Animals in Behavioural Research and Teaching [45] and the legal requirements of the country (France) in which the work was conducted (IBISA platform, agreement F91.114.1, DDPP Essonne, France). All procedures to breed mouse lemurs are conducted in accordance with the European Communities Council Directive (86/609/EEC) and are authorized by the Departmental Veterinary Services (Directive 2010/63/UE). Specifically, for this arboreal primate, housing conditions included cages equipped with branches, various supports, devices to stimulate foraging, and many nesting boxes allowing the animals to express their entire behavioural repertoire. The health and the well-being of captive animals were daily checked by the animal care keepers and a veterinarian. Lastly, only animals that were found naturally dead were used in this study.

## RESULTS

### Maternal characteristics

The naturally dead descendants issued from 637 different mothers aging when giving birth from 1.5 to 8.9 years (mean age 2.2 ± 0.05 yrs). 887 pregnancies (from 1 to 8) were recorded. Owing to breeding protocols in the colony, a majority of pregnancies (63.6%) originated from primiparous females (mean age 1.5 ± 0.05 yrs, N = 564) versus multiparous females (mean age 3.5 ± 0.04 yrs, N = 323). Litter sizes varied from 1 (16.2%) to 2 (48.6%) and 3 offspring (35.2%). Primiparous females produced significantly less number of triplets than multiparous females (29% versus 46%, df_1_, X_2_ = 12.7 – P < 0.001). Pregnancy duration (mean 61.7 ± 0.06 days, N = 815) positively correlated to litter’s size (r = 0.084, P = 0.017, N = 815) but was independent from the mother’s parity or age (df_1/813_, F = 0.09, P = 0.76 and r = 0.044, P = 0.20, N = 815 respectively). The litter masses were evidently highly correlated to the size of the litter (r = 0.757, P < 0.001).

For all pregnancies, BM*oe* (mean 91 ± 0.5g) was positively correlated to mother’ age (r = 0.353 P < 0.001, N = 887) with accordingly significant difference between primiparous and multiparous females (df_1/885_, F = 121, P < 0.001). But age of the mother had no effect on litter masses (r = 0.002, P = 0.95, N 881). Likewise, BM*oe* had no effect on litter masses when using litter size as covariate (df_1/878_, F =1.07, P = 0.30, litter size F = 48.4, P < 0.001).

In conclusion, as their age increased when becoming pregnant, mothers became heavier and produced more numerous offspring, mainly triplets.

### Characteristics of early parameters

#### Birth weight and growth

BM*birth* of the 1365 offspring varied from less than 4g to 10g and differed significantly according to sexes: males (µ = 6.65 ± 0.04 g, N = 643) were heavier than females (µ = 6.47 ± 0.05 g, N = 715, df_1/1356_, F = 7.09, P = 0.008). Their BM*30d* reached on average 33 ± 0.2 g without significant difference between sexes (df_1/1364_, F = 0.03, P = 0.85, N = 1357).

Growth rate averaged 0.88 ± 0.02 g/day without difference between sexes (df_1/1355_, F =0.03, P = 0.8, N = 1357). When 60 days old, young weighted on average 52 ± 0.2 g (N = 1353) with BM*60d* of female offspring significantly higher than that of males (df_1/1352_, F=15.8, P < 0.001 – Fig. 1).

**Fig. 1.**
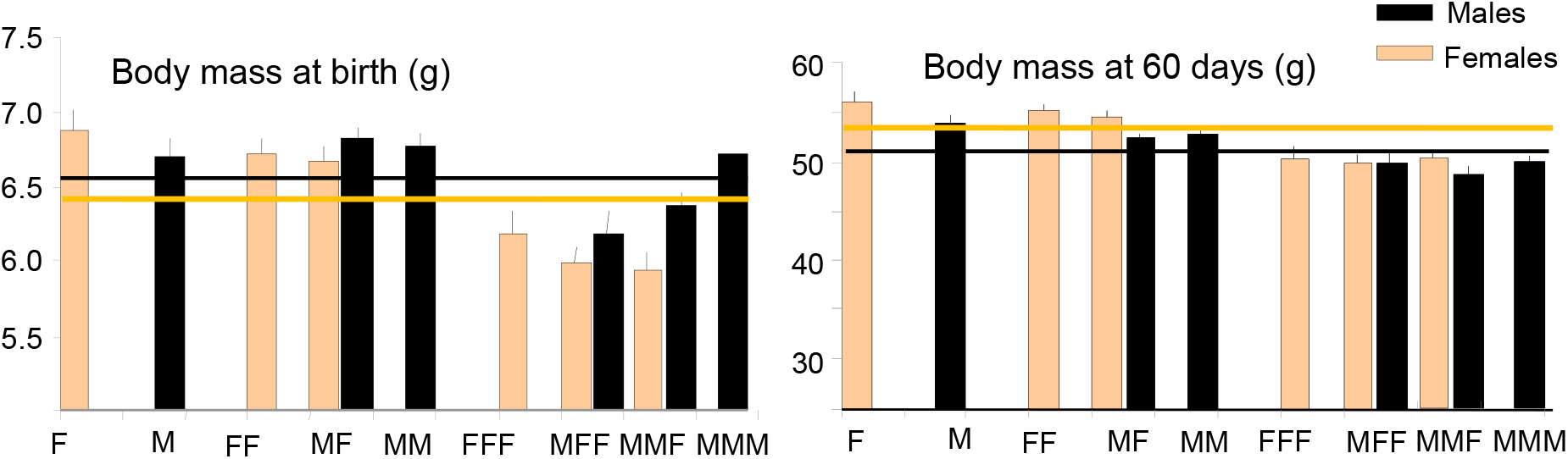
Body mass at birth and at 60 days (g, mean ± sem) of offspring according to the litter composition in which they were born. Mean of males and females were indicated by lines.

**Fig. 2.**
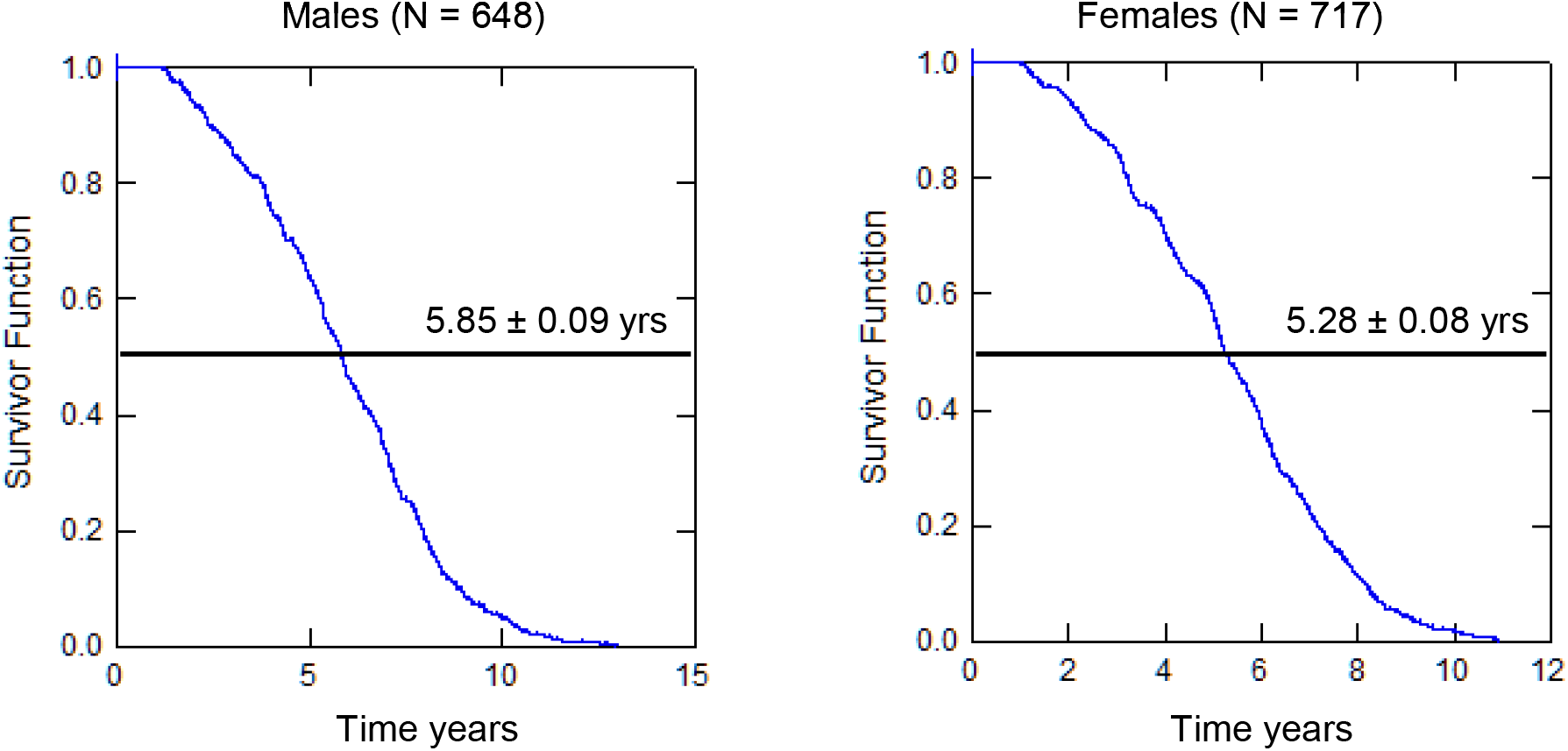
Survival curves of adults with indication of the 50% survival

#### Size and composition of the litter

Litter size affected all parameters (Fig.1). Significant decreases were observed as litter size increased for BM*birth* (df_2/1356_, F=35.4, P < 0.001), growth rate (df_2/1354_, F = 69.2, P < 0.001), BM*30d* (df_2/1354_, F = 83.5, P <0.001) and BM*60d* (df_2/1351_, F=50.7, P < 0.001) in a similar way in both sexes (from P = 0.2 to P = 0.8).

Different effects of litter composition were observed according to their size (Fig. 1). For litters of 1 or 2 babies, BM*birth* were similar between males and females (P = 0.2 and P = 0.3 respectively). By contrast, in triplets, BM*birth* of females were significantly lower compared to those of males (df_1/517_, F = 14.9, P < 0.001). Moreover, BM*birth* in both sexes were significantly lower (df_1/517_, F= 7.56, P = 0.006) in mixed-sex litters (MFF, MMF) compared to those in single-sex litters (FFF, MMM).

With regard to growth rate and BM*30d*, litter composition had no effect in both males and females when using the litter size as covariate (df_/8/1348_, F= 1.04, P = 0.4 and df_8/1347_, F=1.02, P = 0.4, respectively). By contrast, BM*60d* depended to both size (df_2/1351_, F= 50.7, P = 0.001) and composition of the litter (df_8/1344_, F=2.49, P = 0.01 with size as covariate– Fig. 1). Lastly, BM*60d* of males were significantly lower in mixed-sex litters, independently of their size (df_1/639_, F= 10.8, P = 0.001).

### Maternal effects on early parameters

BM*birth* were negatively correlated to the mother’s age in males (r = - 0.095, P = 0.026, N= 643) and in females (r = - 0.105, P = 0.005, N = 715) with a more pronounced effect for females born in triplets (r = -.0.1270, P = 0.036, N =270). By contrast, no significant effect of parity on BM*birth* was observed (df_1/1356_, F = 0.9 P = 0.34). The mother’s age had no effect on growth rate and BM*30d* whatever the sex of offspring (P = 0.4) or the litter’ size (P = 0.8). Likewise, maternal MB*oe* did not affect BM*birth* of both sexes (P = 0.4). By contrast, growth rate and BM*30d* in female offspring were correlated to maternal BM*oe* (r = 0.097, P = 0.009 and r = 0.081, P = 0.03 respectively). More, BM*60d* of female offspring was negatively correlated to the mother’s age (r = - 0.088, P = 0.02, N = 713).

### Early parameters and reproductive success of female offspring

The distribution of successful or unsuccessful females was independent of mother parity (df_2_, X^2^ = 0.1, NS), size (df_2,_ X^2^ = 1.1? NS), composition (df_5_, X^2^ = 13, NS), type (df_1_, X^2^ = 0.5, NS) of the litter in which females were born. BM*birth* and BM*30d* of successful females did not differ from those of unsuccessful females. (P = 0.2 and P = 0.1, respectively). By contrast, BM*60d* of successful females were significantly higher than that of unsuccessful females (df_1/711_, F =6.12, P = 0.014).

#### Early parameters and survival

Using Cox proportional hazards models, the 50% survival of the studied adult population (Fig.2) reached 5.55 ± 0.06 yrs. A highly significant difference was seen between males: µ =5.85 ± 0.09 yrs, N = 648, maximum 12.9 yrs, and females: mean 5.28 ± 0.08 yrs, N = 717, maximum 10.9 yrs (df_1/1356_, F = 21.7, P < 0.001), independent of the size or composition of the litter. Whatever a sex difference in longevity, the shape of survival curves showed no trend difference between sexes.

For female’s offspring, BM*birth* (r = 0.077, P = 0.039, N =715) and BM*30d* (r = 0.090, P = 0.016, N = 714) were positively correlated with survival. By contrast, for male’s offspring, none of these parameters correlated with survival. But in both sexes, a higher BM*60d* correlated with a higher longevity (males r = 0.089, P = 0.024, N = 641 and females, r = 0.080, P = 0.033, N = 713). Lastly, the survival of successfully reproductive females reached 5.60 ± 0.1 yrs, a value significantly higher than those of unsuccessful females (4.61 ± 0.1 yrs, df_1/715_, F = 30.8, P < 0.001).

Characteristics of the mother [body mass prior pregnancy (r = - 0.007, P = 0.8), parity (df_1/1363_, F = 0.03, P = 0.8) or age at conception (r = 0.024, P = 0.38)] had no direct relationship with offspring survival, independently of sexes. Both male and female survival was not linked to litter size (r = - 0.026, P = 0.35). However, a slight effect of the composition of the litter was observed (df_8/1356_, F = 1.09, P = 0.056) – Fig. 3).

**Fig. 3.**
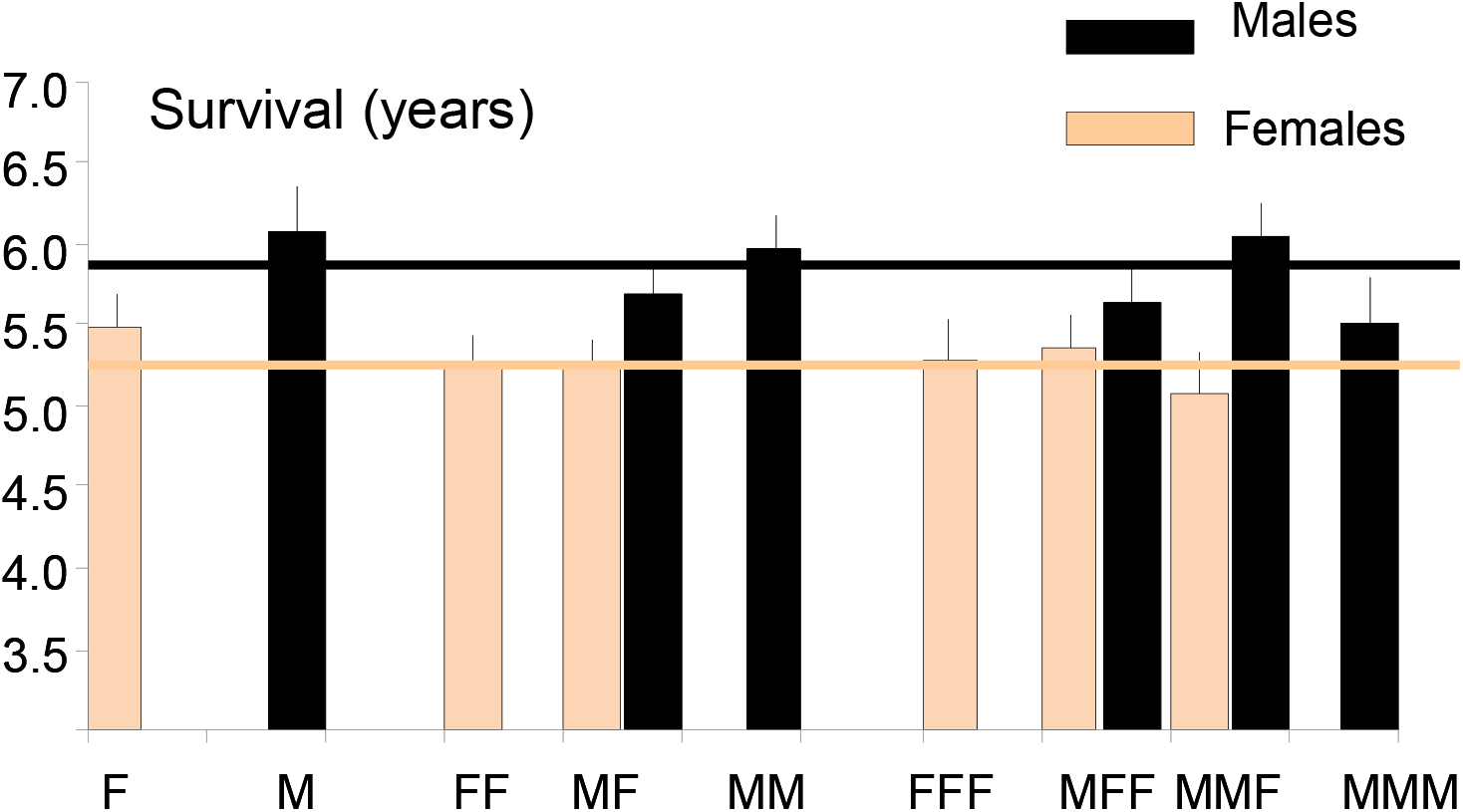
Survival (years) of adults according to the litter composition in which they were born. Mean survivals of males and females were indicated by lines.

#### Pathology and survival

Most of adult males and females died during the LD period (N = 830 in LD versus N = 535 in SD, X^2^ = 63.3, P < 0.001).

The causes of death were determined for only 67% of adults (Table I). For 30% of them, a loss of body mass and asthenia preceded death, whatever the age or pathological syndrome. For animals dying after asthenia without pathology (age = 6.1 ± 0.2 yrs, body mass = 65 ± 0.6 g, N = 249), their BM*60d* were significantly correlated to their longevity (r = 0.164, P = 0.01). More, a significant relationship was found between their longevity and that of their mothers (r = 0.151, N = 235 P = 0.02).

**Table I.** Main causes of deaths (%) according to age’s categories.

| Main causes of deaths | $\leq 3$ yrs<br>N = 139 | 4 - 6 yrs<br>N = 496 | $\geq 7$ yrs<br>N = 276 | $\mu$ age (yrs)<br>± sem |
| --- | --- | --- | --- | --- |
| Kidney failure | 0 | 6.9 | **26.8** | **7.7 ± 0.2** |
| Tumours | 6.5 | 10.1 | **15.9** | **6.6 ± 0.2** |
| Injuries | **34.5** | **31.7** | 6.5 | 4.6 ± 0.1 |
| Reproduction | **24.5** | **19.6** | 2.9 | 4.4 ± 0.1 |
| Others | 9.4 | 9.7 | 9.8 | 5.7 ± 0.2 |

In oldest animals (from 7 to 12.9 yrs, N = 276, mean age = 8.7 ± 0.07 yrs), asthenia without pathology as well as kidney failures could be considered as natural ageing and occurred without differences between males and females independently of the photoperiod. However, in younger animals (mean age 5.9 ± 0.1 yrs), kidney failures were also found, suggesting that this pathology was different from ageing in this case. Tumours (mainly sarcoma) increased with ageing (Table I) and were unrelated to early environment parameters.

A high number of adult animals died after injuries during the breeding season (with or without asthenia or infection). Significant sex differences were observed: - male deaths following sexual competition were different from females (X^2^ = 6.9, P < 0.01, N = 139), and - females dying from injuries in unisex groups were more frequent than males (X^2^ = 9.1, P < 0.01, N = 223). Because these pathologies were related to social interactions, they were independent of early parameters to which the animals have been exposed.

## Discussion

In constant conditions of captivity, where resources are considered as constantly optimal, we characterized early parameters to which mouse lemur’s offspring were exposed and we tested whether these parameters had long term effects on survivorship. Litter size (1 to 3) and composition (sex-mixed or not) allowed to test maternal sex-specific allocation and sibling competition. As expected, values of BM*birth* differed between sexes and were negatively correlated to both litter size and composition. Indeed, the presence of a brother or a sister in the litter had negative effect on BM*birth* of the opposite sex.

At birth, males were significantly heavier than females and this difference remained until weaning, i.e 30-40 days. Thereafter, when the offspring fed themselves, females became heavier than males, owing to their greater ability to contest for food in large litters. As reported across a wide range of taxa, as well as under high nutritional conditions, this strategy of catch-up growth could have long lasting effects on female metabolism rate or insulin regulation in adulthood [15,46,47] or even reduce survival. The occurrence of obesity in some female mouse lemurs could be related to this compensatory growth strategy after weaning.

Mother body condition before pregnancy had no impact on offspring early parameters but the age at conception negatively affected body mass at birth for all offspring, with a higher effect on triplets. Previous analyses on a subset of data have already shown that there was no cost of reproduction for captive female mouse lemurs [49]. In primates, it is generally accepted that older mothers will raise their offspring better than younger mothers, but in captivity, it is not a rule [50]. In mouse lemurs, no reproductive senescence was observed and maternal allocation, through growth rate values, was similar for both sexes regardless of litter size and mother’s age.

When three months old, offspring were grouped by sex and exposed to short photoperiod leading to seasonal fattening. Having a high body mass at that time may facilitate competition for food and for better adaptation to social behaviours. However, none of the offspring’ parameters before weaning correlated with survival. This suggested that sibling competition in large litters, and especially in triplets, could play a major role in the acquisition of a heavy body mass after weaning, as exemplified by its significant relationship with litter composition in both sexes Several studies have demonstrated the important role of litter composition on adult behavioral phenotypes including anxiety, emotional, exploratory or aggressive behaviors [6,7, 51-53]. In mouse lemurs, individual personality and exploratory behaviour have been also related to early parameters [54].

In both sexes, adult survival was highly correlated with body mass after weaning. As exemplified in other vertebrate species, including primates and humans, body mass acquired early in life appears to be a determining factor in survival [55-57].

The survivorship of male mouse lemur is higher than females one. It is well known that males and females differ in longevity in most wild populations [58]. The costs associated with intra-sexual competition in polygynous systems are considered to cause a faster senescence in males. Despite the fact that the mating system of male mouse lemur is related to intra-sexual competition for oestrous females [59], the lifespan of captive males is higher than that of females in captivity. Indeed, a reversed situation was observed in the wild, with female’s survival outliving male’s survival [60]. In captivity, mothers favour their male pups [61], and female offspring were more affected than male offspring by the characteristics of both their mother (age and BM*oe*) and litter composition (mixed-sex litters). In addition to the potential effect of the catch-up growth strategy observed in female offspring, the sex-specific maternal behaviour could have a long-lasting detrimental effect on behavioural or physiological components of females impacting survival. Lastly, the longer life expectancy of captive males could be linked to the fact that sexual competition is limited to very short periods each breeding season (less than 4 weeks).

A high BM60*d* also appeared to play a determining role in females’ reproductive success, fitting with a silver-spoon effect. Indeed, successfully reproductive females have a BM*60d* and a survival significantly higher than that of unsuccessful females. Successful reproduction was thus consistently associated to increased longevity in mouse lemurs. The higher survival in females that have successfully raised offspring could be attributed to the protective effect of reproduction against oxidative stress, as demonstrated in several species [62-65].

More, BM60*d* as well as other parameters of primiparous mothers correlated positively with early parameters of their offspring. This suggested a potential intergenerational effect, referring to a transmission of environmental effects on mother to their developing offspring [66,67].

Even if early environment conditions can play an important role in captive mouse lemur’s survival and also in reproductive success of both sexes, according to litter composition [40-41), cumulative effects of genetics and of adult environmental conditions may strongly interact to affect fitness and survival [14]. Genetically, offspring survival is partially dependent on the longevity of their parents [68], especially that of their mothers. Indeed, for oldest animals, the maternal longevity positively correlated with their longevity, as exemplified in other captive primate’s species [69].

Adult environment plays a major role on adult survival. In captivity, mouse lemurs are maintained in small groups and, especially during the breeding season, aggressive behaviours lead to severe injuries, with consequently reduced survival in young and adult animals during sexual competition. Aggressive behaviours between adult females are responsible for numerous deaths, more numerous than in males. These behaviours could explain the reduced survival of captive females and could be related to the unusual situation of Malagasy prosimians where “female dominance” seems the rule [70]. In captive males, deaths following injuries are also consequences of sexual competition under constraint captive conditions. Other causes of deaths such as kidney failures or tumours observed in oldest animals can be attributed to normal ageing, as observed in another mouse lemur’s colony [71]. But, kidney failures in younger animals are a known consequence of social stress induced by captivity [72]. All effects of intra- or inter-sexes social interactions during adulthood may outweigh the effect of favourable early environment and would therefore minimize the silver-spoon effect on captive mouse lemur’s survival. Quantifying the effects of early-life conditions on adult survival in the wild is challenging owing to multiple environmental factors, especially resource availability and predation. However, perhaps through differences in personality, individuals of both sexes that have a high body mass after weaning have greater potential for survival through better adaptation to constraints either social in captivity or environmental in the wild. Future research is needed to determine to what extent early conditions have long-term effects on behavioural or physiological adulthood phenotype.

## Acknowledgements

We are extremely indebted to I. Hardy for creating the “*Mouse lemur life history traits*” and A. Anzeraey for continuing to complete the database (1995 - present). We would like to thank the animal care keepers, L. Dezaire, S. Gondor and I. Hiron and E. Pasquier for their contribution to the excellent care of the mouse lemurs.

